# Above- and below-ground trait coordination across 90 angiosperm and gymnosperm tree species

**DOI:** 10.1101/2025.03.05.641585

**Authors:** Anvar Sanaei, Karl Andraczek, Lena Kretz, Florian Schnabel, Ronny Richter, Anja Kahl, Nicole Nabel, Lea von Sivers, Tom Künne, Julia Leonore van Braak, Ronja Felicitas Hofmann, Carolin Sophie Hensel, Karin Mora, Hannes Feilhauer, Christian Wirth, Alexandra Weigelt

**Affiliations:** Institute of Biology, Leipzig University, 04103 Leipzig, Germany; German Centre for Integrative Biodiversity Research (iDiv) Halle-Jena-Leipzig, 04103 Leipzig, Germany; Leibnitz Centre for Agricultural Landscape Research (ZALF), Müncheberg, Germany; Department Conservation Biology and Social-Ecological Systems, Helmholtz Centre for Environmental, Research (UFZ), Leipzig, Germany; Chair of Silviculture, Faculty of Environment and Natural Resources, University of Freiburg, Freiburg, Germany; Remote Sensing Center for Earth System Research, Department of Remote Sensing in Geo-Ecosystem research, Leipzig University, 04103, Leipzig, Germany; Max-Planck-Institute for Biogeochemistry, 07745 Jena, Germany

**Keywords:** acquisitive strategy, collaboration gradient, conservative strategy, fine-root traits, leaf economics spectrum, leaf traits, plant economics spectrum, root economics space

## Abstract

Quantifying the variation in plant traits reveals the trade-offs involved in plant ecological strategies and is fundamental to understanding underlying plant fitness mechanisms. Thus, the ecological success of plant species in a certain habitat may depend on the coordinated performance of both leaves and roots. However, despite the growing interest in trait variation, it is still uncertain i) to what extent the leaf economics spectrum (LES) and root economics space (RES) hold across locally coexisting tree species and ii) whether leaf and fine-root traits are correlated. In a research arboretum, we simultaneously measured eight key traits in leaves and fine-roots on 270 individuals belonging to 90 tree species, encompassing both angiosperm and gymnosperm species. We find varied plant resource strategies associated with leaves and fine-roots for angiosperms and gymnosperms. We observe a clear LES for gymnosperms and a clear RES for angiosperms. Our results support the existence of a correlation between analogous leaf and fine-root traits across all species. However, varying trait coordination across clades indicates varying resource acquisition strategies above- and belowground, highlighting the need to consider large-scale phylogenetic relatedness to better understand plant fitness.

## 1. Introduction

One of the main long-standing challenges in ecology is to elucidate plant community assembly mechanisms (Grime, 1977; Westoby & Wright, 2006; Lavorel & Grigulis, 2012). Plant traits provide a comprehensive mechanistic approach to understanding how plant species exist and survive in a given habitat (Lavorel & Garnier, 2002). Leaf and root traits are vital for plant fitness maintenance and have significant contributions to biogeochemical cycles (Wright et al., 2004; Weigelt et al., 2021). Coordination of leaf and fine-roots is essential for the effective use of limited resources, thereby optimizing plant growth and survival (Reich, 2014). This synchronized functioning of leaves and fine-roots should encourage some coordination or trade-off between leaf and fine-root traits. To date, researchers have been exploring i) if the relationships between leaf or fine-root traits identified on a global scale are consistent at a local scale, and ii) if leaf and fine-root traits are coordinated and thus represent the same trait syndromes above- and below-ground. Three main trait syndromes, focusing on different plant organs, have been described, namely the leaf economics spectrum (LES), the root economics space (RES), and the whole plant economics spectrum (PES), which we will introduce in the following paragraphs.

At the leaf level, a continuum between resource acquisition and conservation based on the potential for a return on investment of carbon is depicted as the LES (Wright *et al*., 2004; Reich, 2014). Following this, fast-growing and short-lived species are characterized by low leaf mass per area, leaf dry matter content, thinner leaves, and high leaf nitrogen concentration, resulting in a higher growth rate (Wright *et al*., 2004; Shipley *et al*., 2006). On the other hand, slow-growing species tend to have opposing leaf traits, leading to a lower growth rate but a higher lifespan (Wright *et al*., 2004; Shipley *et al*., 2006). These trade-offs have been observed among plant communities (Wright *et al*., 2004; Reich, 2014), across various growth forms (Shiklomanov *et al*., 2020) and phylogenetic clades (Maynard *et al*., 2022) at the global scale. Evidence from previous studies suggests that conservative and acquisitive plant strategies are conserved within plant clades. For instance, angiosperms seem to be more dominated by acquisitive strategies, while gymnosperms are more dominated by conservative strategies (Givnish, 2002; Maynard *et al*., 2022; Ribeiro *et al*., 2022). Indeed, gymnosperms typically possess a longer leaf lifespan than angiosperms and conserve more biomass to sustain photosynthesis (Berendse & Scheffer, 2009). In contrast, angiosperms offset a short leaf lifespan by increasing productivity at the leaf level, resulting in rapid resource returns to maintain efficient photosynthesis (Givnish, 2002; Wright *et al*., 2004). A recent global study indicated that angiosperms are indeed more acquisitive than gymnosperms, characterized by higher leaf nitrogen concentration while having a low leaf mass per area and leaf thickness (Maynard *et al*., 2022). Importantly, while the LES concept has greatly advanced our ability to understand leaf trait variations, it has rarely been tested for multiple species at the local scale.

In contrast to leaves, roots do not only serve the absorption of multiple resources but also fulfil other functions such as providing structural support and interaction with soil microbial communities (Kramer-Walter *et al*., 2016; Kong *et al*., 2019; Bergmann *et al*., 2020), all while encountering complex biotic and abiotic constraints (Valverde-Barrantes *et al*., 2016; Weemstra *et al*., 2016). Consequently, complementary to the one-dimensional LES, fine-roots have been depicted as multidimensional based on a global dataset of 1,810 plant species, known as a root economics space (RES; Bergmann et al., 2020). Under this premise, the widely accepted economics spectrum for leaves is transferable to fine-roots, i.e., roots less than two mm in diameter (Reich, 2014; Roumet *et al*., 2016; Kramer-Walter *et al*., 2016). This so-called conservation gradient in roots is manifested by high root nitrogen concentrations as an acquisitive strategy and low root tissue density as a conservative strategy (Fig. 1; Kramer-Walter et al., 2016; Bergmann et al., 2020). Indeed, this gradient depicts a continuum from less metabolically active tissues with high root tissue density to highly metabolically active tissues with high root nitrogen concentrations, reflecting a quick return on investment (Wright et al. 2004; Reich 2014). However, in contrast to leaves, interactions with arbuscular mycorrhizal fungi (AMF), which support plants’ ability to acquire nutrients and water, have a significant impact on below-ground resource uptake strategies (Weemstra *et al*., 2016; McCormack & Iversen, 2019) and related traits that form a ‘collaboration’ gradient. Thus, the collaboration gradient is on the one side characterized by high root diameter, associated with high mycorrhizal colonization and an ‘outsourcing’ strategy of resource acquisition and on the other side with thin roots with high specific root length that follow a ‘do-it-yourself’ strategy (Bergmann *et al*., 2020). This implies that species with thinner roots maximize root surface for independently exploring the soil for resources, while those with thicker roots are more reliant on mycorrhizal for nutrient acquisition (Bergmann *et al*., 2020). Similar to leaf traits, gymnosperms and angiosperms differ substantially in root morphology, architecture, anatomy, and nutrient concentrations, leading to distinct responses to their environment, and possibly, clade-specific coordination of root traits (Zhang *et al*., 2017). Thus, similar to leaf traits above-ground, below-ground angiosperms are more dominated by thinner roots with more fine-root tips and higher root nitrogen concentration (Valverde-Barrantes *et al*., 2016; Wang *et al*., 2019), whereas gymnosperms are more dominated by thicker roots and use a more conservative strategy (Ma *et al*., 2018; Langguth *et al*., 2024). As such, less ramified fine-roots and slower root proliferation in gymnosperms (Bauhus & Messier, 1999; Withington *et al*., 2006) result in higher dependence on mycorrhizal for soil resource acquisition (Liu *et al*., 2015). While gymnosperms had been underrepresented, more recently, Langguth et al. (2024) identified a RES for 34 gymnosperm species growing under three different site conditions in the USA and Poland.

**Figure 1.**
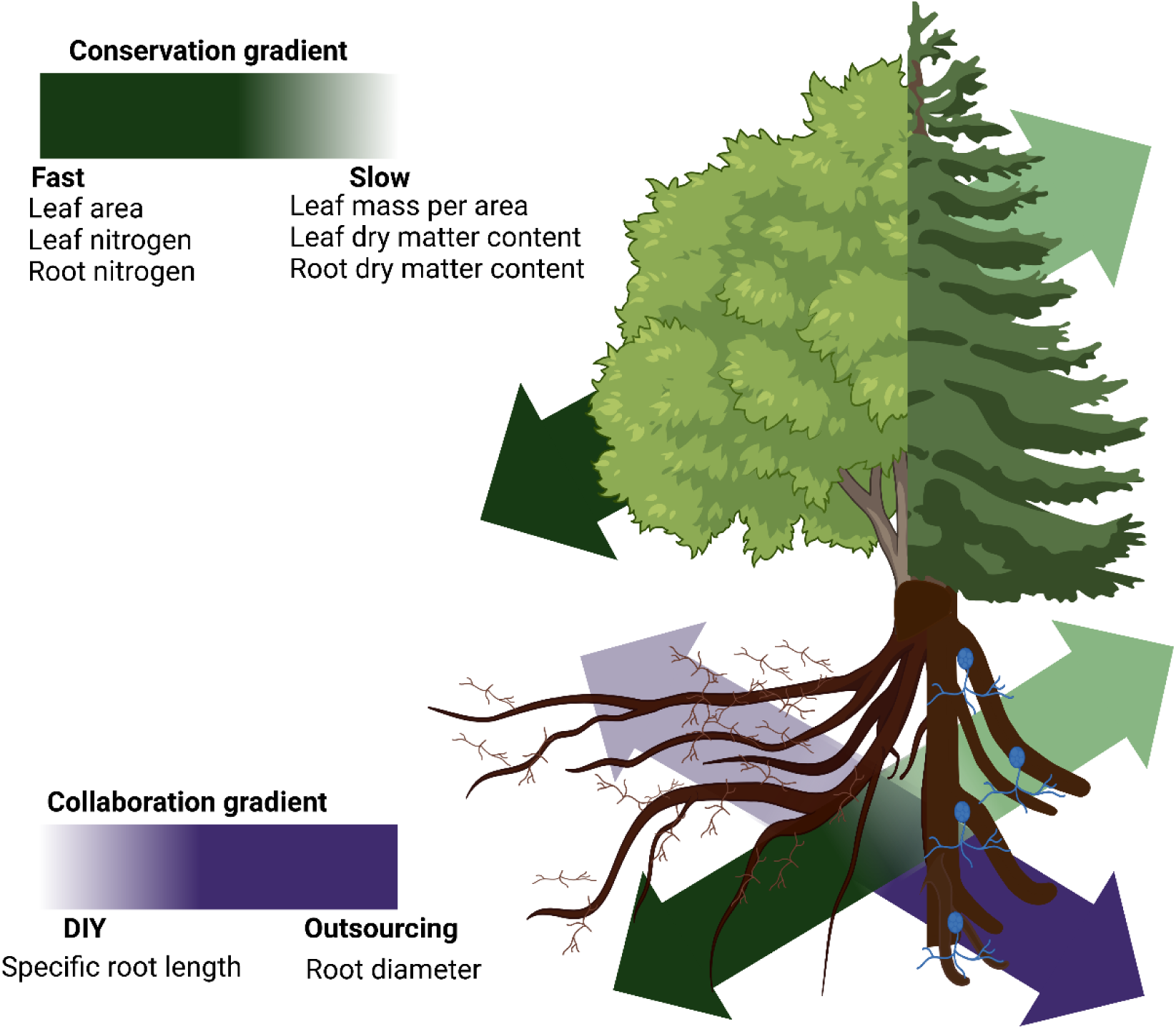
Conceptual framework illustrating the leaf economics spectrum (LES), the root economics space (RES), and the plant economics spectrum (PES). The diagram is inspired by Weigelt et al. (2021). The figure was created with BioRender.com.

Lastly, above- and below-ground organs do not exist in isolation but rather exhibit diverse levels of structural and functional linkages, resulting in the ability to perform multiple functions. Following this rationale, the plant economics spectrum (PES) incorporates multiple functional traits from different plant organs, e.g., leaf, stem and fine-roots, that are involved in resource uptake and coordinates them along a single fast-slow spectrum (Reich, 2014; Burton *et al*., 2020; Weigelt *et al*., 2021; Li *et al*., 2022). Specifically, based on the leaf and fine-root fast-slow continuum, plant species with a conservative resource strategy in leaves are supposed to have a conservative resource strategy in roots. Thus, higher leaf dry matter content and leaf mass per area, but low leaf nitrogen concentrations, should be mimicked by a conservative resource strategy in fine-root as well, i.e., higher root tissue density and root dry matter content but low root nitrogen concentrations (Fig. 1). Similarly, acquisitive resource strategies in leaves and fine-roots should be aligned as well (Reich, 2014; Weigelt *et al*., 2021). However, the root collaboration gradient is thought to be orthogonal to leaf and root conservation gradients (Fig. 1; Weigelt et al., 2021; Zhang et al., 2024). While the PES provides a more mechanistic understanding of overall plant strategies in terms of growth, reproduction, and survival (Reich, 2014; Díaz *et al*., 2016; Weigelt *et al*., 2021), it is still not clear if and to what extent analogous leaf and fine-root traits are correlated. Observations at both global and local scales report ambiguous relationships; while some studies identified a strong coupling of leaf and fine-root traits (Weigelt *et al*., 2021; Mueller *et al*., 2024), others found a decoupled relationship (Burton *et al*., 2020; Carmona *et al*., 2021). These contrasting results are likely driven by differences in trait and species selection (Weigelt et al., 2023, but see Bueno et al., 2023), trait plasticity, and the variation in biotic and abiotic environments (Weigelt *et al*., 2021). However, the mechanisms underlying these patterns remain largely unresolved.

Although the global-scale reports for the LES (Wright *et al*., 2004; Díaz *et al*., 2016) and RES relationships (Bergmann *et al*., 2020; Weigelt *et al*., 2021) are remarkably consistent, findings at finer spatial and taxonomic scales have reported mixed results (Messier et al., 2017a,b; Anderegg et al., 2018). Hence, the question is: Why did previous studies, at different scales, observe such varied patterns in leaf and fine-root traits? A possibility is that, at the global scale, environmental gradients, e.g., climatic and soil properties, act as the main drivers of plant functional trait variation (Freschet *et al*., 2017; Joswig *et al*., 2022). In contrast, locally coexisting tree species may exhibit local adaptations to specific biotic and abiotic conditions, which may shape distinct trade-offs, resulting in varying trait dimensions depending on the strength of the trade-offs (Messier *et al*., 2017a; Defrenne *et al*., 2019; Yang & Russo, 2024). Another possible explanation for the discrepancies between global and local trait dimensions is that trait covariation may be weaker in fine-scale studies, likely due to the limited number of investigated species (Anderegg *et al*., 2018).

Our study focused on quantifying the variations between a suite of key functional traits in the leaves and fine-roots of 90 tree species, including 65 angiosperms and 25 gymnosperms, growing under the same environmental conditions in a research arboretum in Germany. This unique set-up allowed us to examine the correspondence of LES, RES and PES in the absence of confounding effects of environmental gradients on a fine spatial scale. Specifically, we hypothesized (H1) that leaf trait coordination of the investigated 90 tree species is consistent with the global trait framework of the LES (Fig. 1). Second, we hypothesized (H2) that fine-root trait coordination of the 90 tree species is aligned with the trait framework of the RES (Fig. 1). Finally, we incorporated leaf and fine-root traits to quantify a comprehensive PES. We hypothesized (H3) that pairs of potentially analogous leaf and fine-root traits, defined as a conservation gradient, reveal the same syndromes and that conservation gradients are independent of the root collaboration gradient. Yet, because angiosperms and gymnosperms are known to differ considerably in evolutionary history and ecological strategies, we expected different growth strategies and associated differences in leaf and fine-root traits, leading to dissimilar patterns in LES, RES and PES (Fig. 1). We therefore examined trait space correlation for all species and separately for both clades.

## 2. Materials and Methods

### 2.1 Study area and experimental design

The study was conducted at the research Arboretum ARBOfun located in Großpösna, Germany (51°16′ N and 12°30′ E). Between 2012 and 2014, five individuals of 100 tree species, chosen for their ecological and economic value for European forests, were planted with 5.8 m inter-distance in a randomized block design. However, due to mortality, some species are no longer alive, and not all species have five replicates available. The average annual temperature of the arboretum is approximately 9.7 °C and the average annual precipitation is around 520 mm (DWD Climate Data Center [CDC], Station Leipzig/Halle, ID 2932). The soil type is a Luvisol, with a pH of 5.7 (Ferlian *et al*., 2017). For the present study, we chose 270 tree individuals of 90 species (three replicates for each species; Table S1; Fig. 2).

**Figure 2.**
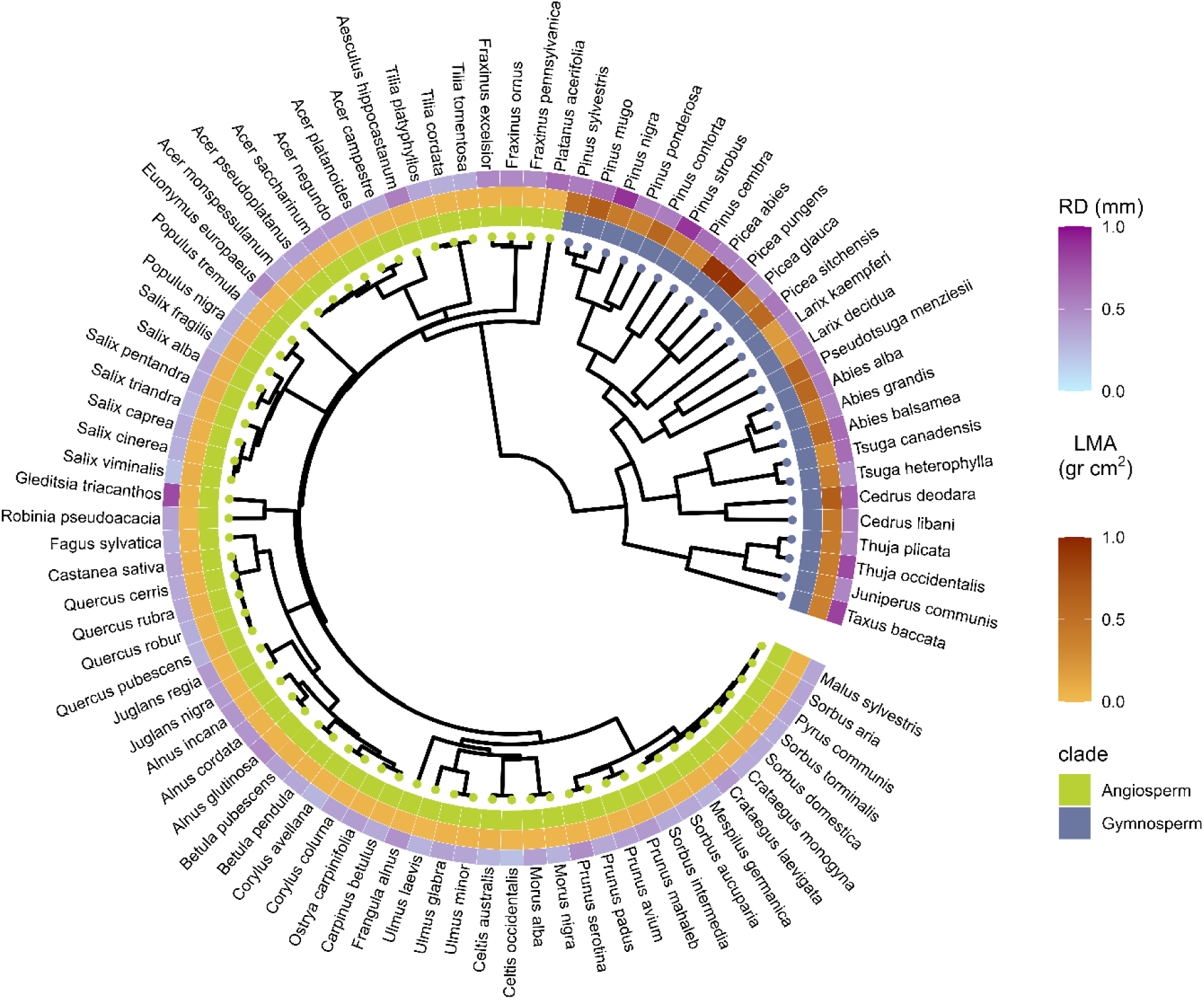
Phylogenetic tree of 90 tree species used in this study. The inner colors for the Latin names represent different clades, and the outer colors show two traits, namely root diameter (RD) and leaf mass per area (LMA), involved in trait interplay that are mapped onto each species.

### 2.2. Leaf sampling and measurement

Following standard protocols (Cornelissen *et al*., 2003), at the end of June 2023, fully expanded and intact sun-exposed leaves—at least five leaves for angiosperms and 40 needles for gymnosperms—were randomly selected and collected from each individual tree. The collected leaves (including both lamina and petiole) were scanned at 600 dpi with a flatbed scanner (Epson Expression 11000XL, UK), and the images were analyzed for leaf area using WinFolia (Regent Instrument, Canada). The scanned leaves were weighed to obtain their fresh weights. Subsequently, the samples were oven-dried at 60 °C for five days and reweighed to obtain the dry weight. The leaf mass per area (LMA) was computed by dividing the dry mass of the leaves by their total fresh area. The mean leaf area per leaf (LA) was computed by dividing the total leaf area by the number of leaves. The leaf dry matter content (LDMC) was determined by dividing the mean leaf dry weight by the mean leaf fresh weight. Finally, leaf nitrogen concentration (LN, % of dry weight) was measured using an elemental analyzer (VarioEL II, Elementar).

### 2.3. Root sampling and measurement

After loosening the soil around the target tree using a digging fork, the roots were uncovered carefully by hand. If a root of higher order was found, it was followed back to the main stem of the target tree to verify its identity. Then intact root branches containing at least the first five-order roots, with the most distal root tip as the first-order root, were collected. The root samples, including adherent soil, were wrapped in moist paper, sealed in plastic bags and stored in a cooling box before being transported to the laboratory where they were stored for maximum 24 h at 4 °C. After washing root samples, the sample of each individual tree was divided into three sub-samples: one for examining morphological traits, one for examining architectural traits, and one as backup sample. For morphological trait examination, we first identified the first three root orders under a stereo microscope, and then a flatbed scanner (Epson Expression 11000XL, UK) was used to scan the fine-root samples at 600 dpi resolution, where the fine-roots were arranged in trays without any overlap. After scanning, the root pieces were collected, and weighed to get fresh weight, then oven-dried at 60 °C for >48 h, weighed to determine dry weight, and ground for chemical analysis.

We analyzed morphological root traits at individual tree level based on the root scans using the RhizoVision Explorer version 2.0.3 (Seethepalli *et al*., 2021). The average root diameter (RD) was obtained using the scanned images with RhizoVision. We then calculated specific root length (SRL) as the ratio between total root length and root dry weight. Next, root dry matter content (RDMC) was calculated as the ratio between root fresh weight and dry weight. Root nitrogen concentration (RN, % of dry weight) was measured using an elemental analyzer (VarioEL II, Elementar).

### 2.4. Statistical analysis

To identify the main dimensions of trait variation, we performed principal component analyses (PCAs) of leaf and fine-root traits. To do so, the first series of PCAs were performed on leaf traits, including LN, LDMC and LMA for all species consistent with studies investigating fast-slow leaf trait coordination (Díaz *et al*., 2016) as well as individual PCAs for each clade (angiosperms and gymnosperms). We also included LA as it impacts light interception, energy balance, and water regulation (Díaz *et al*., 2016). The second series of PCAs were performed on the root traits, including RN, RDMC, RD and SRL for all species as well as individual PCAs for each clade. Hence, in this second step we focused on those traits that were also analyzed in recent studies investigating the root economics space (Bergmann *et al*., 2020; Weigelt *et al*., 2021). However, rather than using RTD, we used RDMC, as our goal was to have an analogous root trait to LDMC. In order to verify this selection, we found a highly significant relationship between RTD and RDMC (*R^2^* = 0.43, *P* < 0.001), consistent with a previous study (Birouste *et al*., 2014). Finally, the third series of PCAs were extended to the whole set of leaf and fine-root traits by integrating fine-root and leaf traits. In this step, we concentrated on leaf and fine-root traits that play key roles in plant form and function (Carmona *et al*., 2021; Weigelt *et al*., 2021). To account for potential phylogenetic dependency among species, we used phylogenetically informed PCA for the analyses across all species and each clade based on species mean trait values. For that, based on the available backbone phylogeny (Zanne *et al*., 2014), we constructed a phylogenetic tree using the *phylomatic* function of the ‘Branching’ package (Torres-Jimenez, 2024). We then visualized the phylogenetic tree for all 90 species, annotating clades and two functional traits (LMA and RD) using the *ggtree* and *gheatmap* functions of the ‘ggtree’ package (Yu *et al*., 2017). Then, the phylogenetic signals for all leaf and fine-root traits were estimated using the Blomberg’s K test and computed using the *phylosig* function in the ‘phytool’ package (Revell, 2012). To complement the results, we subsequently performed bivariate ordinary least squares (OLS) regression using the ‘stats’ package and phylogenetic generalized least squares (PGLS) regression using the function of *pgls* in the ‘caper’ package (Orme *et al*., 2023). To meet the linear regression assumptions, all traits were log-transformed before performing pairwise bivariate regression. Lastly, the phylogenetically informed PCAs were conducted using the *phyl.pca* function of the ‘phytool’ package (Revell, 2012) on log-transformed trait data. All the statistical analyses were conducted using the R v.4.3.2 statistical environment (R Core Team, 2024).

## 3. Results

### 3.1. Phylogenetic conservatism and bivariate relationships among traits

Across the 90 species, all leaf and fine-root traits, except LDMC, showed significant phylogenetic signals, as evidenced by Blomberg’s *K* (Table 1). The phylogenetic signals showed different patterns when we looked at each clade; for angiosperms, only RD and for gymnosperms, RN showed significant phylogenetic signals (*P* = 0.002 and 0.023, respectively; Table S2), while SRL and LA showed marginally significant phylogenetic signals for angiosperms and gymnosperms, respectively (*P* = 0.058 and 0.076, respectively; Table S2). Furthermore, using species trait means, we found a significant difference between clades (angiosperms and gymnosperms) for all leaf and fine-root traits except for RDMC (Fig. S1).

**Table 1.** Variation in eight leaf and fine-root traits measured in 90 tree species. Phylogenetic signal and significance were tested using a two-sided Blomberg’s K test. SD is standard deviation, CV is coefficient of variation, Max is maximum and Min is minimum.

| Trait | Abbreviations | Unit | Mean | SD | CV | Max | Min | Blomberg's *K* | *P* value |
| --- | --- | --- | --- | --- | --- | --- | --- | --- | --- |
| Leaf nitrogen | LN | % | 1.799 | 0.522 | 0.290 | 3.200 | 0.953 | **0.037** | **0.037** |
| Leaf dry matter content | LDMC | g g<sup>-1</sup> | 0.418 | 0.035 | 0.085 | 0.491 | 0.353 | 0.029 | 0.169 |
| Leaf area | LA | cm<sup>2</sup> | 41.514 | 67.737 | 1.632 | 354.826 | 0.131 | **0.086** | **0.001** |
| Leaf mass per area | LMA | g cm<sup>-2</sup> | 0.168 | 0.216 | 1.284 | 0.927 | 0.031 | **0.073** | **0.002** |
| Specific root length | SRL | mm mg<sup>-1</sup> | 33.600 | 19.576 | 0.583 | 117.493 | 6.792 | **0.054** | **0.005** |
| Root diameter | RD | mm | 0.440 | 0.141 | 0.321 | 0.885 | 0.227 | **0.052** | **0.004** |
| Root dry matter content | RDMC | g g<sup>-1</sup> | 0.340 | 0.057 | 0.168 | 0.502 | 0.215 | **0.041** | **0.020** |
| Root nitrogen | RN | % | 1.382 | 0.496 | 0.359 | 3.481 | 0.588 | **0.054** | **0.005** |

The pairwise PGLS bivariate regression among leaf and fine-root traits for all tree species showed that LN was negatively correlated with LDMC, LA, and LMA, while it was significantly positively correlated with RN (Fig. 3). LDMC was significantly positively correlated with both LMA and RDMC, while it was negatively correlated with RN. LMA was negatively correlated with RN (Fig. 3). Moreover, SRL was significantly negatively correlated with both RD and RDMC, while it was significantly positively correlated with RN. RN was negatively correlated with both RDMC and RD (Fig. 3). Consistent with the results for all species, the bivariate PGLS regressions among leaf traits for angiosperms showed a negative correlation between LA and LN, as well as a positive correlation between LDMC and RDMC (Fig. S2). LDMC was negatively correlated with RD. Similar to all species, angiosperm SRL was negatively correlated with RD, while it was positively correlated with RN. RDMC was negatively correlated with RN (Fig. S2). In contrast to all species, RD and LDMC were negatively correlated (Fig. S2). In contrast with angiosperms, LN in gymnosperms was negatively correlated with both LMA and LDMC, but also showed a positive correlation with RN (Fig. S3). In addition, LMA was positively correlated with LDMC but negatively correlated with RN (Fig. S3). In line with results for all species, SRL showed a negative correlation with RD and RDMC as well as a positive correlation with RN (Fig. S3).

**Figure 3.**
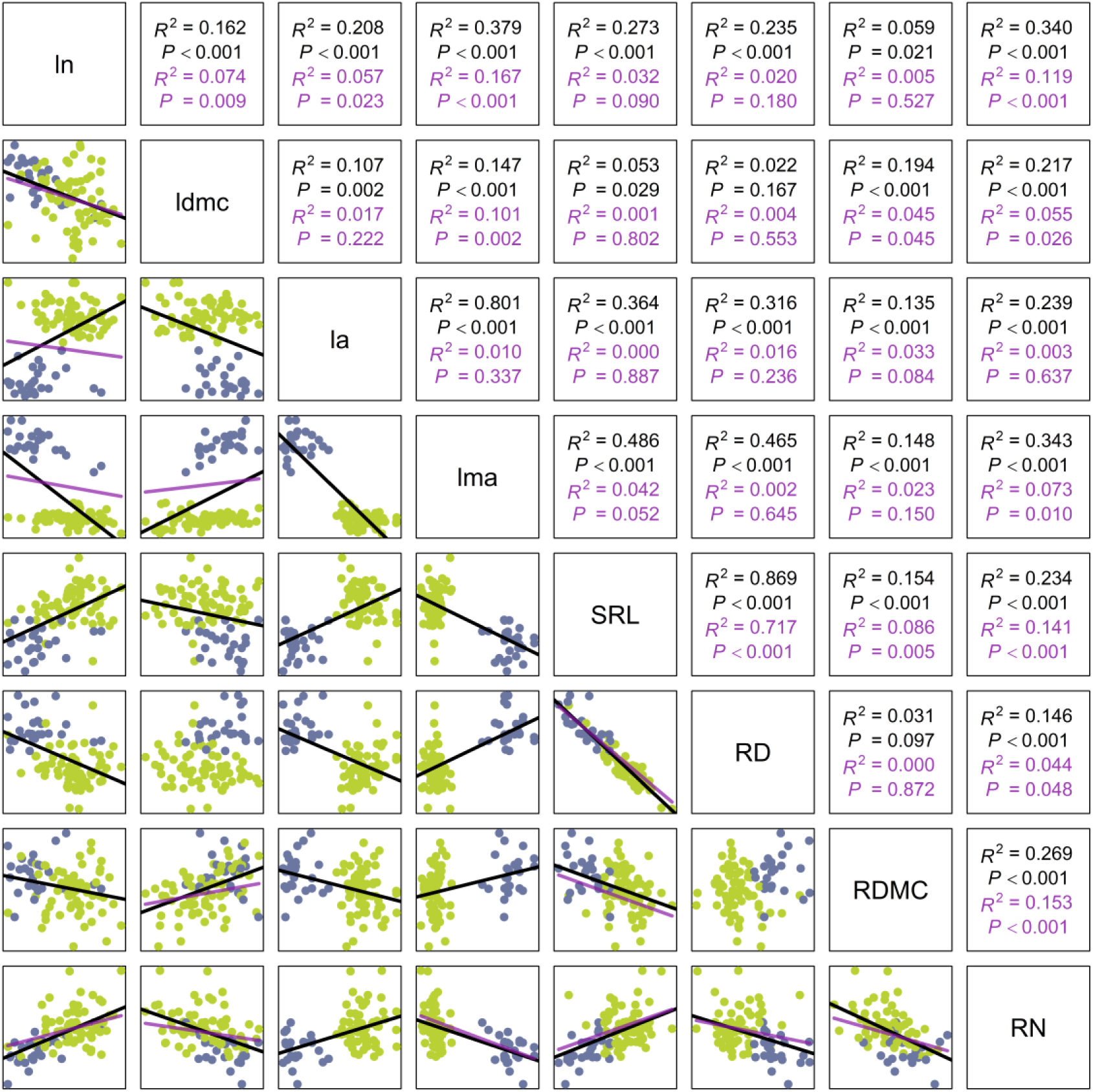
Pairwise correlations among leaf and fine-root traits for all 90 tree species, including angiosperms (green) and gymnosperms (violet). The bivariate relationships among leaf and fine-root traits for all tree species are assessed using ordinary least squares (OLS) and phylogenetic generalized least squares (PGLS) models. Significant correlations are presented by regression lines colored black for OLS models and pink for PGLS models. ln is leaf nitrogen concentration; ldmc is leaf dry matter content; la is leaf area; lma is leaf mass per area; SRL is specific root length; RD is average root diameter; RDMC is root dry matter content and RN is root nitrogen concentration.

### 3.2. Multivariate leaf and fine-root trait coordination

The leaf trait coordination for all species showed that the first two PC axes together accounted for 69% of the total variance (Fig. 4a). The first PC axis captured the variation in LMA and LDMC, with gymnosperms characterized by higher LMA and LDMC (Figs. 4a, S4a; Table S3). The second PC axis captured a gradient from LN to LA. We found a comparable pattern of leaf trait coordination for angiosperms to that observed in all species, with a shift in LMA and LDMC from the first PC axis to the second PC axis and also LA and LN shifting from the second PC axis to the first PC axis (Figs. 4b, S4b). These first two PC axes together explained 55% of the variance (Fig. 4b; Table S3). Gymnosperm leaf trait pattern showed that the two PC axes together explained 79% of leaf trait variation (Fig. 4c; Table S3). Here, the first PC axis captured the acquisitive-conservative leaf strategy with LMA and LDMC on the one and LN on the other side. In addition, LA, as a size-related leaf trait, was loaded on the second PC axis (Figs. 4c, S4c; Table S3).

**Figure 4.**
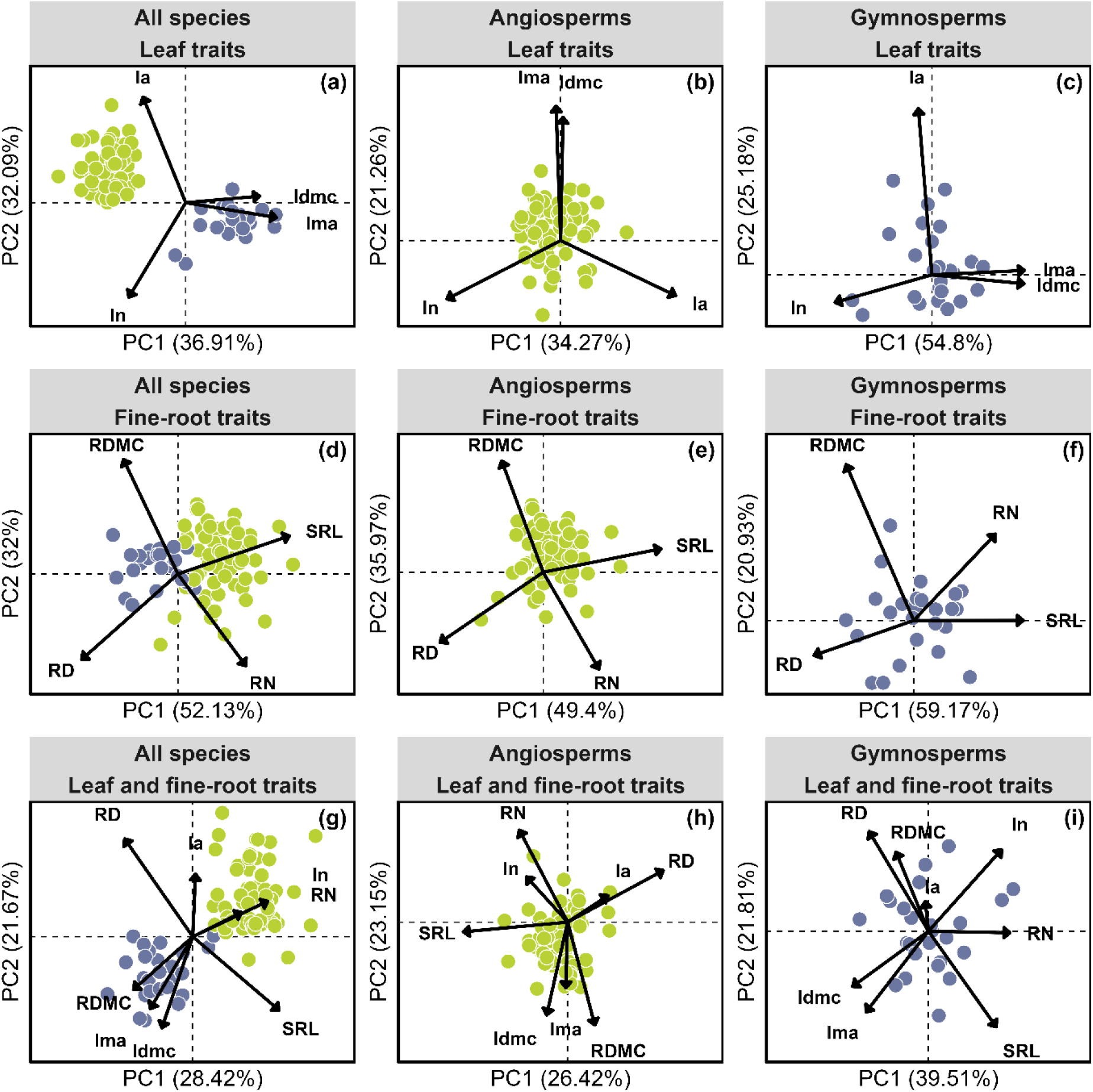
Phylogenetically informed principal component analyses of tree species based on species’ mean trait values for (a, d, g) all species (*n* = 90); (b, e, h) angiosperm species (green, *n* = 65); and (c, f, i) gymnosperm species (violet, *n* = 25). Trait abbreviations follow the caption in Figure 3. See Supporting Information Table S3 and Figure S4 for the loading scores.

Consistent with other studies on the RES, the multivariate fine-root pattern across all species was depicted by two separate dimensions (Fig. 4d), which cumulatively explained 84% of the variation. In particular, the first PC axis captured the variation in SRL and RD, representing the root collaboration gradient (Figs. 4d, S4d; Table S3), where gymnosperms show a higher RD than angiosperms. The second PC axis captured the variation in RN and RDMC, showing the root conservation gradient (Figs. 4d, S4d; Table S3). We found the same directionality of fine-root trait pattern among angiosperms, as observed for all species, with the first two PC axes accounting for 85% of the variation (Fig. 4e). In particular, the first PC axis captured the variation in RD and SRL, while the second PC axis captured the variation in RDMC and RN (Figs. 4e, S4e; Table S3), indicating an independent depiction of the root collaboration and conservation gradients. For gymnosperms, the first two PC axes of fine-root traits captured 80% of the variation (Fig. 4f). Unlike the fine-root trait coordination for all species and angiosperms, the RN of gymnosperms did not show a clear trade-off with the RDMC, but instead a positive correlation with SRL, thus being loaded on the first PC axis alongside SRL, while RDMC was loaded on the second PC axis (Figs. 4f, S4f; Table S3).

The PCA for the full set of leaf and fine-root traits showed that the first two PC axes together explained 50% of the variance (Fig. 4g; Table S3).The first PC axis captured the variation in RN, SRL and RDMC, while the second PC axis captured the variation in LDMC, and RD (Figs. 4g, S4g; Table S3), again showing gymnosperms represented by a higher LDMC and RDMC but lower SRL and RN (Fig. 4g; Table S3). We found that RD was loaded, with more or less similar loading, onto both the first and second PC axes (Figs. 4g, S4g; Table S3). We observed a comparable trait coordination pattern of the whole set of leaf and fine-root traits for angiosperms to that depicted in all species, with a shift in RN and RDMC from the first axis to the second axis. In particular, the first two PC axes together explained 49% of the variability; the first PC axis captured the variation in RD and SRL, showing an independent root collaboration gradient (Figs. 4h, S4h; Table S3). The second PC axis captured the variation in LDMC, RDMC and RN, showing conservation gradients of fine-root traits (Figs. 4h, S4h; Table S3). Finally, the whole set of leaf and fine-root traits for gymnosperms showed that the majority of the trait variation (61%) was captured by the first two PC axes. Unlike the trait coordination observed in all species and angiosperms, there were some trait-trait relationships shifting in gymnosperms (Fig. 4i). In particular, LDMC was decoupled from RDMC, resulting in LDMC negatively loading on the first PC axis alongside LMA, LN, RN and SRL, while RD, and SRL, to a lesser extent, loaded on the second PC axis as well (Figs. 4i, S4i; Table S3).

## 4. Discussion

Considering eight leaf and fine-root functional traits that are crucial for ecological strategies, resource acquisition and plant fitness, we explored if general directions of the LES and RES manifest across 90 tree species. We found distinct life history trade-offs associated with above- and below-ground plant organs for angiosperm and gymnosperm species. Gymnosperms showed an opposing pattern to angiosperms, consistent with previous studies (Maynard *et al*., 2022), most likely due to their distinct resource uptake strategies. Our study further demonstrates a clear LES primarily for gymnosperms, while a clear RES exists across all species and for angiosperms. Finally, by integrating leaf and fine-root traits, we found distinct relationships between the corresponding above- and below-ground traits in angiosperms and gymnosperms. Specifically, we found that the conservative ends of the leaf and root conservation gradients were strongly correlated in angiosperms and all species; conversely, the acquisitive ends of the conservation gradients showed stronger correlation in gymnosperms and all species. The root collaboration gradient, an axis that was much clearer in angiosperms than in gymnosperms, formed an orthogonal axis to leaf and root conservation gradients, consistent with recent studies conducted at both local (Han *et al*., 2022) and global scales (Bergmann *et al*., 2020; Weigelt *et al*., 2021). Overall, the varied integration of above- and below-ground patterns within the two clades highlights the importance of clade-specific analyses in improving our understanding of plant resources and, ultimately, plant fitness.

### 4.1. Leaf trait coordination

The rate of plant photosynthesis and consequently plant growth is primarily related to LN and LMA, along a single fast-slow gradient (Wright *et al*., 2004; Reich, 2014). Considering these leaf traits and LA, we found a clear separation between angiosperm and gymnosperm species, where gymnosperms loaded on the “conservative” side and angiosperms loaded on the “acquisitive” side, supporting our first hypothesis (H1). Indeed, this difference in leaf traits among gymnosperm and angiosperm species reflects their contrasting adaptation to seasonality, environmental stress, defense against herbivory, and efficiency in resource investment (Wright *et al*., 2004; Díaz *et al*., 2016). In particular, the trade-off between carbon gain and leaf lifespan has been attributed to LMA (Díaz *et al*., 2016), where a higher LMA, as observed here for gymnosperms, is associated with leaves being thick with higher physical strength and a longer lifespan (Onoda *et al*., 2017). In addition, the size-related trait LA is associated with light interception efficiency and leaf thermodynamics, and the construction cost per leaf area of larger LA leaves, here for angiosperms, is mostly higher than that of leaves with a smaller area (Milla & Reich, 2007; Wright *et al*., 2017). We argue that the separation between angiosperm and gymnosperm species is partly due to the greater differences in leaf traits between the two clades; as such, we found a higher LN and larger LA in angiosperms, while the larger LMA and higher LDMC in gymnosperms (Fig. S1). Our results align with previous findings that highlight the significance of LMA in separating between angiosperms and gymnosperms (Stahl *et al*., 2013). However, we did not observe a clear fast-slow continuum defined in the LES as a gradient from LN to LMA and LDMC along the same axis. Nevertheless, a significant negative correlation (indicating a possible ‘trade-off’) was found for the pairwise relationships between LN with both LMA and LDMC (Fig. 3).

Given the changing trade-off between traits across clades, separate multivariate analyses of leaf traits in angiosperms and gymnosperms showed different coordination patterns, in line with our expectations. Leaf trait variation in gymnosperms was depicted by two axes; the first axis reflects fast-slow resource strategy and was represented by LMA, LDMC, and LN, and the second axis was represented by LA. The first axis supports a persistence-performance trade-off of the LES (Wright *et al*., 2004; Gomarasca *et al*., 2023). On the one end, leaves invest more nitrogen in persistence to extend their lifespan and longevity; on the other end, they invest more nitrogen in building RuBisCO to enhance photosynthetic capacity (Luo *et al*., 2021). For angiosperms, however, a trade-off between LA and LN was observed, this was also supported by significant negative phylogenetic relationships between LA and LN (Fig. S3). Overall, considering the observed leaf trait patterns, the LES was much clearer in gymnosperms. In a global study, however, the general LES held true for both angiosperms and gymnosperms (Maynard *et al*., 2022). This disagreement might be partly attributed to the number of included species, i.e., local vs. global, and also the context dependency of the LES, where leaf trait dimensions on a global and local scale might not be necessarily consistent (Messier *et al*., 2017b). Another related point is that, Maynard et al. (2022) included many other traits, such as leaf thickness, tree height, crown diameter and height and stem diameter, which might lead to altering the overall trait patterns (Weigelt *et al*., 2023). Altogether, inconsistent direction of the LES across angiosperm and gymnosperm species suggests that leaf traits are not driven by the same processes in these clades and highlights their distinct resource-use strategies, adaptation to environmental drivers, different photosynthesis rates, and the growing season (Stahl *et al*., 2013; Augusto *et al*., 2015). Thus, trait-based approaches to reveal mechanistic insight on trait-function relationships might benefit from clade-specific analyses.

### 4.2. Fine-root trait coordination

Like most studies on root traits, we found multidimensionality in fine-root traits, as evidenced by two independent dimensions for all species, i.e., including angiosperms and gymnosperms (Bergmann *et al*., 2020; Weigelt *et al*., 2021), supporting our second hypothesis (H2). The first dimension reflects the dependency of species on mycorrhiza for resource uptake, manifested by trade-offs between the two fine-root traits SRL and RD, i.e., the root collaboration gradient (Bergmann *et al*., 2020). The other dimension is represented by trade-offs between fast vs. slow resource return on investment, known as the root conservation gradients (Bergmann *et al*., 2020). Within the root collaboration axis, angiosperm species were separated on the ‘do-it-yourself’ side with a higher SRL, while gymnosperms were separated on the ‘outsourcing’ side with a higher RD. This finding supports previous studies demonstrating varying mycorrhizal dependency of gymnosperms and angiosperms for nutrient acquisition, with gymnosperms having thicker roots that are more reliant on mycorrhizal association, while angiosperms have thinner roots that allow them to explore the soil more efficiently and independently for resources (Valverde-Barrantes *et al*., 2016; Ma *et al*., 2018; Wang *et al*., 2019).

Consistent with our expectations, we found differences in the RES between the two clades. For angiosperms, similar to all species, we found a clear RES where fine-root trait coordination is reflected by two distinct axes: the first axis represents the root collaboration gradient, and the second axis represents the conservation gradient. This pattern is consistent with the recently published RES framework (Bergmann *et al*., 2020) and with a more recent local study for angiosperms (Wang *et al*., 2024). For gymnosperms, we did not observe a clear RES, because RN was unrelated to RDMC, positively correlated with SRL, and thus opposed to RD. However, a more recent study found that the RES holds true across 34 gymnosperm species (of which only 12 species were included in our analysis, Fig. 2; Table S1) using the first-order root traits (Langguth *et al*., 2024), partly contrasting our findings. This suggests that including a more complete set of gymnosperm species in our analysis might have revealed a RES. A distinct trait coordination between angiosperms and gymnosperms could be explained by the unique characteristics of the two clades: First, a unique carbon allocation strategy where angiosperms are often fast-growing, allocating more carbon to their leaves and reproductive organs, whereas gymnosperms allocate more carbon to structural organs, enhancing their lifespan. Second, while the RES was primarily established based on AMF symbiosis (Bergmann *et al*., 2020), the varied types of mycorrhizal symbiosis species between angiosperms and gymnosperms might be another factor influencing the observed differences in fine-root traits between the two clades. As such, strong differences in root traits, distinctions in life-history strategy, as well as distinct nutrient sources in the RES between AMF and ectomycorrhizal (EMF) species have been reported (Brundrett & Tedersoo, 2018; Averill *et al*., 2019; Yan *et al*., 2022). More specifically, out of 25 gymnosperms, only 4 species were associated with AMF, and the remaining 21 species were associated with EMF. Conversely, most of the angiosperms (38 species; Table S1) were associated with AMF. In line with this, for angiosperms, *Robinia pseudoacacia* L. from family Fabaceae has the potential for nitrogen fixing and showed the highest RN; also, the Alnus genera formed N-fixing nodules, which might contribute to soil nutrient acquisition. For example, more recently, a higher SRL, RN and AMF colonization has been found for N-fixing species compared to not N-fixing species (Marcellus *et al*., 2024). Overall, these differences between angiosperms and gymnosperms might lead to the distinct RES pattern, further highlighting distinct below-ground constraints and strategies.

### 4.3. Leaf and fine-root trait coordination

By integrating leaves and fine-roots of 90 species to test the PES concept, we found a more complex trait space than that discovered by the small set of traits considered in LES or RES. As such, the two PC axes accounted for only 50% of the total variance, indicating that the overall plant form and function are shaped by a much more complex trait space. Specifically, traits were coordinated primarily along four dimensions. The first PC axis is mostly related to SRL alongside the root conservation gradient ascribed by RN and RDMC. However, RD was related to the second and first axes (loadings: 0.67 and -0.63, respectively), alongside LDMC, representing the conservative end of the conservation gradient in leaves. The third axis relates mostly to LN and LA. However, we found that analogous leaf and fine-root traits were located on the same side of the PCA, yet they were not loaded on the same axis (Figs. 4, S4; Table S3). Nonetheless, they showed significant correlation, as evidenced by pairwise correlation, with a stronger correlation between LN and RN compared to LDMC and RDMC (Fig. 3). In agreement with our results, the stronger bivariate relationships have been reported for chemical traits (RN and LN) than for morphological traits, e.g., LDMC and RDMC (Valverde-Barrantes *et al*., 2015; Mueller *et al*., 2024). Partially supporting our third hypothesis (H3), we did observe a collaboration gradient where RD and SRL were clearly negatively correlated, albeit not fully loaded on one axis of the PCA (Figs. 3, S4; Table S2). Our results do not fully match a most recent global study, which found that the conservation gradients in leaves and fine-roots were independent of the collaboration gradient (Weigelt *et al*., 2021). These differences again highlight the distinct drivers shaping global vs. local scale trait coordination, as adaptations to biotic and abiotic conditions on local scales might lead to different combinations of above- and below-ground traits (Messier *et al*., 2017a). The limited number of species in our dataset, compared to more global-scale studies, may have restricted trait variation and prevented us from identifying clear trade-offs between species (Anderegg *et al*., 2018). Furthermore, Weigelt et al. (2021) used the classical definition of root diameter for fine-roots(≤ 2 mm); while this study focused on the first three-order roots. Order-based fine root selection supports the importance of absorptive roots in resource acquisition for tree growth (Sanaei *et al*., 2025), and has been shown to affect RN, SRL and RTD (McCormack *et al*., 2015). Consistent with the global study, our results further showed that LA, a size-related above-ground trait, was not related to the LES or RES (Figs. 3, 4; Díaz et al., 2016). On a local scale, LA variation might partially be attributable to local biotic and abiotic conditions (Zhou *et al*., 2022), while on a global scale, it might be driven by elevational and latitudinal gradients (Wright *et al*., 2017).

Next, we explored the degree to which analogous leaf and fine-root traits, defined by conservation gradients in both leaves and fine-roots, are consistent across the two clades. Trait coordination patterns differed between angiosperms and gymnosperms, as expected, due to changes in above- and below-ground trait-trait relationships. In particular, in angiosperms, the first axis represents a clear independent root collaboration gradient (SRL-RD) and the second axis shows the conservation gradient (RDMC-RN) (Figs. 4, S2; Table S3). In addition, in angiosperms, there was a correlation between LDMC and RDMC but not between LN and RN; this was also evidenced by the pairwise correlations (Fig. S2). However, in gymnosperms, we found a correlation between the analogous chemical leaf and fine-root traits (LN-RN) but no correlation between the analogous morphological leaf and fine-root traits (LDMC-RDMC), as well as the existence of a conservation gradient in leaves (LDMC-LN trade-off) but not in fine-roots. Together, this reveals that some tree species with similar fine-root traits might not necessarily have similar leaf traits, and this varying combination of above- and below-ground might reveal various adaptive phenotypic strategies (Carmona *et al*., 2021). Since functional traits reflect plant responses to environmental changes (Lavorel & Garnier, 2002), uncorrelated fine-root and leaf traits may indicate that above- and below-ground traits are not exactly driven by similar biotic and abiotic factors, or the degree to which these factors impact them may differ (Carmona *et al*., 2021; Luo *et al*., 2024). Moreover, the lack of strong coordination between morphological leaf and fine-root traits in gymnosperms might be due to more dissimilar development of leaves and fine-roots compared to angiosperms. More specifically, in gymnosperms, needle-shaped leaves have slow turn-over, but angiosperm leaves have shorter life spans. We might argue that, in angiosperms, when leaves are at the conservative end of the conservation gradients (higher LDMC), fine-roots show the same strategy (higher RDMC); in contrast, for gymnosperms, when leaves are at the acquisitive end of the conservation gradient (higher LN), fine-roots also show the same strategy (higher RN). Lastly, in gymnosperms both SRL and RD similarly loaded onto both the first and second PC axes. We still observed that the collaboration gradient in gymnosperms is orthogonal to above- and below-ground conservation traits; however, this collaboration axis was less clear than that of angiosperms. This might result from the narrower range of gymnosperm species in our analysis, belonging to only three families (Pinaceae with 22 species, Cupressaceae with two species and Taxaceae with only one species), compared to the broader range of angiosperm families included in this study (Table S1). The varied above- and below-ground alignment between the two clades might help species in acquiring additional resources and ultimately, optimizing fitness. Altogether, the integration of above- and below-ground organs differs between angiosperms and gymnosperms; this reveals a clear distinction in carbon allocation strategies between these two clades, highlighting fundamental differences in their leaf and fine-root systems (Stahl *et al*., 2013; Wang *et al*., 2019).

## Conclusions

Our results, across all species, showed a clear separation between angiosperms and gymnosperms, highlighting different resource strategies associated with leaf and fine-root functional traits. Different trait coordination of the LES or RES was observed between angiosperms and gymnosperms; while a clear LES exists in gymnosperms, a clear RES holds in angiosperms. Collectively, varied trait coordination patterns between the two major clades here reaffirm the idea that distinct plant life history strategies are controlled by unique trade-offs. Thus, conducting separate analyses for the two clades could provide more mechanistic insight on trait-function relationships. Furthermore, understanding local-scale trait variation is crucial for identifying plant strategies that evolve in response to local environmental conditions, which may be obscured in large-scale studies. The selection of the eight fine-root and leaf traits for this study was partly based on their inclusion in previous LES and RES trait analyses and their significance to key ecological strategies. This did not include above- and below-ground size-related traits (Díaz *et al*., 2016; Weigelt *et al*., 2021), some of which may reveal further aspects of tree form and function. Thus, we acknowledge that a deeper understanding of plant form and function and the mechanisms underlying resource acquisition strategies at local scales profits from research combining multiple above- and below-ground traits—morphology-, physiology-, lifespan-, chemistry-, anatomy-, mycorrhizal colonization-, architecture-, hydraulic- and size-related traits—measured across many species.

## Acknowledgments

We thank members of the botanical garden of Leipzig University for the maintenance of the research arboretum and numerous student helpers for support with field and laboratory work. We would like to acknowledge Ines Hilke and the service group ‘RoMA’ at the MPI for Biogeochemistry for measuring C and N concentrations in leaf and root tissues. A.S is supported by the Saxon State Ministry for Science, Culture and Tourism (SMWK) – [3-7304/35/6-2021/48880].

## Author Contributions

A.S., A.W. and C.W. conceived the ideas and developed the concept of the study. A.S., L.K., F.S., R.R., A.K., N.N., L.S., T.K., J.L.B., R.F.H., C.S.H., C.W. and A.W. contributed to data collection. A.S. performed the data analysis and led the writing of the manuscript with support from K.A., F.S., L.K., R.R., K.M., H.F., C.W. and A.W. All authors contributed critically to the drafts and gave final approval for publication.

## Data Availability

We will store the dataset in the public repository once the paper is accepted.

