## Supplementary Information File for "Above- and below-ground trait coordination across 90 angiosperm and gymnosperm tree species"

**Table S1**. List of 90 tree species used in this study. AMF is arbuscular mycorrhizal fungi and EMF is ectomycorrhizal fungi.

| NO. | Species | Family | Order | Clade | Mycorrhizal symbiosis |
|---|---|---|---|---|---|
| 1 | *Acer campestre* L. | Sapindaceae | Sapindales | Angiosperms | AMF |
| 2 | *Acer monspessulanum* L. | Sapindaceae | Sapindales | Angiosperms | AMF |
| 3 | *Acer negundo* L. | Sapindaceae | Sapindales | Angiosperms | AMF |
| 4 | *Acer platanoides* L. | Sapindaceae | Sapindales | Angiosperms | AMF |
| 5 | *Acer pseudoplatanus* L. | Sapindaceae | Sapindales | Angiosperms | AMF |
| 6 | *Acer saccharinum* L. | Sapindaceae | Sapindales | Angiosperms | AMF |
| 7 | *Aesculus hippocastanum* L. | Sapindaceae | Sapindales | Angiosperms | AMF |
| 8 | *Alnus cordata* (Lois.) Desf. | Betulaceae | Fagales | Angiosperms | EMF |
| 9 | *Alnus glutinosa* Medik. | Betulaceae | Fagales | Angiosperms | EMF |
| 10 | *Alnus incana* L. | Betulaceae | Fagales | Angiosperms | EMF |
| 11 | *Betula pendula* | Betulaceae | Fagales | Angiosperms | EMF |
| 12 | *Betula pubescens* | Betulaceae | Fagales | Angiosperms | EMF |
| 13 | *Carpinus betulus* L. | Betulaceae | Fagales | Angiosperms | EMF |
| 14 | *Castanea sativa* Mill. | Fagaceae | Fagales | Angiosperms | EMF |
| 15 | *Celtis australis* L. | Cannabaceae | Rosales | Angiosperms | AMF |
| 16 | *Celtis occidentalis* L. | Cannabaceae | Rosales | Angiosperms | AMF |
| 17 | *Corylus avellana* L. | Betulaceae | Fagales | Angiosperms | EMF |
| 18 | *Corylus colurna* L. | Betulaceae | Fagales | Angiosperms | EMF |
| 19 | *Crataegus laevigata* (Poir.) DC. | Rosaceae | Rosales | Angiosperms | AMF |
| 20 | *Crataegus monogyna* Jacq. | Rosaceae | Rosales | Angiosperms | AMF |
| 21 | *Euonymus europaeus* L. | Celastraceae | Celastrales | Angiosperms | AMF |
| 22 | *Fagus sylvatica* L. | Fagaceae | Fagales | Angiosperms | EMF |
| 23 | *Frangula alnus* L. | Rhamnaceae | Rosales | Angiosperms | AMF |
| 24 | *Fraxinus excelsior* L. | Oleaceae | Lamiales | Angiosperms | AMF |
| 25 | *Fraxinus ornus* L. | Oleaceae | Lamiales | Angiosperms | AMF |
| 26 | *Fraxinus pennsylvanica* L. | Oleaceae | Lamiales | Angiosperms | AMF |
| 27 | *Gleditsia triacanthos* L. | Fabaceae | Fabales | Angiosperms | AMF |
| 28 | *Juglans nigra* L. | Juglandaceae | Fagales | Angiosperms | AMF |
| 29 | *Juglans regia* L. | Juglandaceae | Fagales | Angiosperms | AMF |
| 30 | *Malus sylvestris* (L.) Mill | Rosaceae | Rosales | Angiosperms | AMF |
| 31 | *Mespilus germanica* L. | Rosaceae | Rosales | Angiosperms | AMF |
| 32 | *Morus alba* L. | Moraceae | Rosales | Angiosperms | AMF |
| 33 | *Morus nigra* L. | Moraceae | Rosales | Angiosperms | AMF |
| 34 | *Ostrya carpinifolia* Scop. | Betulaceae | Fagales | Angiosperms | EMF |
| 35 | *Platanus acerifolia* (Aiton) Willd. | Platanaceae | Proteales | Angiosperms | AMF |
| 36 | *Populus nigra* L. | Salicaceae | Malpighiales | Angiosperms | EMF |
| 37 | *Populus tremula* L. | Salicaceae | Malpighiales | Angiosperms | EMF |
| 38 | *Prunus avium* L. | Rosaceae | Rosales | Angiosperms | AMF |
| 39 | *Prunus mahaleb* L. | Rosaceae | Rosales | Angiosperms | AMF |
| 40 | *Prunus padus* L. | Rosaceae | Rosales | Angiosperms | AMF |
| 41 | *Prunus serotina* Ehrh. | Rosaceae | Rosales | Angiosperms | AMF |
| 42 | *Pyrus communis* L. | Rosaceae | Rosales | Angiosperms | AMF |
| 43 | *Quercus cerris* L. | Fagaceae | Fagales | Angiosperms | EMF |
| 44 | *Quercus pubescens* Willd. | Fagaceae | Fagales | Angiosperms | EMF |

| | | | | | |
|---|---|---|---|---|---|
| 45 | *Quercus robur* L. | Fagaceae | Fagales | Angiosperms | EMF |
| 46 | *Quercus rubra* L. | Fagaceae | Fagales | Angiosperms | EMF |
| 47 | *Robinia pseudoacacia* L. | Fabaceae | Fabales | Angiosperms | AMF |
| 48 | *Salix alba* L. | Salicaceae | Malpighiales | Angiosperms | EMF |
| 49 | *Salix caprea* L. | Salicaceae | Malpighiales | Angiosperms | EMF |
| 50 | *Salix cinerea* L. | Salicaceae | Malpighiales | Angiosperms | EMF |
| 51 | *Salix fragilis* L. | Salicaceae | Malpighiales | Angiosperms | EMF |
| 52 | *Salix pentandra* L. | Salicaceae | Malpighiales | Angiosperms | EMF |
| 53 | *Salix triandra* L. | Salicaceae | Malpighiales | Angiosperms | EMF |
| 54 | *Salix viminalis* L. | Salicaceae | Malpighiales | Angiosperms | EMF |
| 55 | *Sorbus aria* | Rosaceae | Rosales | Angiosperms | AMF |
| 56 | *Sorbus aucuparia* | Rosaceae | Rosales | Angiosperms | AMF |
| 57 | *Sorbus domestica* L. | Rosaceae | Rosales | Angiosperms | AMF |
| 58 | *Sorbus intermedia* (Ehrh.) Pers. | Rosaceae | Rosales | Angiosperms | AMF |
| 59 | *Sorbus torminalis* (L.) Crantz | Rosaceae | Rosales | Angiosperms | AMF |
| 60 | *Tilia cordata* Mill. | Malvaceae | Malvales | Angiosperms | EMF |
| 61 | *Tilia platyphyllos* Scop. | Malvaceae | Malvales | Angiosperms | EMF |
| 62 | *Tilia tomentosa* Moench | Malvaceae | Malvales | Angiosperms | EMF |
| 63 | *Ulmus glabra* Huds. | Ulmaceae | Rosales | Angiosperms | AMF |
| 64 | *Ulmus laevis* Pall. | Ulmaceae | Rosales | Angiosperms | AMF |
| 65 | *Ulmus minor* Mill. | Ulmaceae | Rosales | Angiosperms | AMF |
| 66 | *Abies alba* (L.) Mill. | Pinaceae | Pinales | Gymnosperms | EMF |
| 67 | *Abies balsamea* | Pinaceae | Pinales | Gymnosperms | EMF |
| 68 | *Abies grandis* (Douglas ex D. Don) Lindley | Pinaceae | Pinales | Gymnosperms | EMF |
| 69 | *Cedrus deodara* G.Don | Pinaceae | Pinales | Gymnosperms | EMF |
| 70 | *Cedrus libani* A. Rich. | Pinaceae | Pinales | Gymnosperms | EMF |
| 71 | *Juniperus communis* L. | Cupressaceae | Cupressales | Gymnosperms | AMF |
| 72 | *Larix decidua* Mill. | Pinaceae | Pinales | Gymnosperms | EMF |
| 73 | *Larix kaempferi* (Lamb.) Carr. | Pinaceae | Pinales | Gymnosperms | EMF |
| 74 | *Picea abies* (L.) H. Karst. | Pinaceae | Pinales | Gymnosperms | EMF |
| 75 | *Picea glauca* (Moench) Voss | Pinaceae | Pinales | Gymnosperms | EMF |
| 76 | *Picea pungens* Engelm. | Pinaceae | Pinales | Gymnosperms | EMF |
| 77 | *Picea sitchensis* (Bong.) Carr. | Pinaceae | Pinales | Gymnosperms | EMF |
| 78 | *Pinus cembra* L. | Pinaceae | Pinales | Gymnosperms | EMF |
| 79 | *Pinus contorta* Douglas | Pinaceae | Pinales | Gymnosperms | EMF |
| 80 | *Pinus mugo* Turra | Pinaceae | Pinales | Gymnosperms | EMF |
| 81 | *Pinus nigra* J.F. Arnold | Pinaceae | Pinales | Gymnosperms | EMF |
| 82 | *Pinus ponderosa* Douglas ex C. Lawson | Pinaceae | Pinales | Gymnosperms | EMF |
| 83 | *Pinus strobus* L. | Pinaceae | Pinales | Gymnosperms | EMF |
| 84 | *Pinus sylvestris* L. | Pinaceae | Pinales | Gymnosperms | EMF |
| 85 | *Pseudotsuga menziesii* (Mirbel) Franco | Pinaceae | Pinales | Gymnosperms | EMF |
| 86 | *Taxus baccata* L. | Taxaceae | Cupressales | Gymnosperms | AMF |
| 87 | *Thuja occidentalis* L. | Cupressaceae | Cupressales | Gymnosperms | AMF |
| 88 | *Thuja plicata* | Cupressaceae | Cupressales | Gymnosperms | EMF |
| 89 | *Tsuga canadensis* (L.) Carrière | Pinaceae | Pinales | Gymnosperms | EMF |
| 90 | *Tsuga heterophylla* (Raf.) Sarg. | Pinaceae | Pinales | Gymnosperms | EMF |

**Table S2.** Blomberg's K value of leaf and fine-root traits for angiosperm (*n* = 65) and gymnosperm (*n* = 25) species.

| Trait | Abbreviations | Unit | Angiosperms | | Gymnosperms | |
|---|---|---|---|---|---|---|
| | | | Blomberg's *K* | *P* value | Blomberg's *K* | *P* value |
| Leaf nitrogen | ln | % | 0.057 | 0.152 | 0.072 | 0.628 |
| Leaf dry matter content | ldmc | $g\ g^{-1}$ | 0.032 | 0.426 | 0.069 | 0.693 |
| Leaf area | la | $cm^{-2}$ | 0.031 | 0.489 | **0.145** | **0.058** |
| Leaf mass per area | lma | $g\ cm^{-2}$ | 0.025 | 0.638 | 0.085 | 0.429 |
| Specific root length | SRL | $mm\ mg^{-1}$ | **0.075** | **0.076** | 0.055 | 0.884 |
| Root diameter | RD | mm | **0.121** | **0.002** | 0.066 | 0.690 |
| Root dry matter content | RDMC | $g\ g^{-1}$ | 0.021 | 0.670 | 0.101 | 0.254 |
| Root nitrogen | RN | % | 0.014 | 0.839 | **0.164** | **0.023** |

**Table S3.** Results of phylogenetically informed principal component analyses based on the correlation matrix of 90 tree species for all species, angiosperms and gymnosperms, as shown in Figure 4. lma is leaf mass per area; la is leaf area; ln is leaf nitrogen concentration; ldmc is leaf dry matter content; SRL is specific root length; RDMC is root dry matter content; RD is average root diameter; RN is root nitrogen concentration. Displayed data are the variance explained by each principal component (PC) and the loading scores of the leaf and fine-root traits on the first six PC axes.

| **All species (*n* = 90)** | | | | | | | **Angiosperms (*n* = 65)** | | | | | | | **Gymnosperms (*n* = 25)** | | | | | | |
|---|---|---|---|---|---|---|---|---|---|---|---|---|---|---|---|---|---|---|---|---|
| *Leaf traits* | | | | | | | | | | | | | | | | | | | | |
| **Figure 4a** | **PC1** | **PC2** | **PC3** | **PC4** | | | **Figure 4b** | **PC1** | **PC2** | **PC3** | **PC4** | | | **Figure 4c** | **PC1** | **PC2** | **PC3** | **PC4** | | |
| Variance (%) | 0.369 | 0.321 | 0.203 | 0.107 | | | Variance (%) | 0.333 | 0.319 | 0.213 | 0.125 | | | Variance (%) | 0.548 | 0.252 | 0.119 | 0.081 | | |
| eigenvalue | 1.215 | 1.133 | 0.902 | 0.653 | | | eigenvalue | 1.171 | 1.130 | 0.922 | 0.708 | | | eigenvalue | 1.481 | 1.004 | 0.691 | 0.568 | | |
| lma | 0.805 | -0.118 | 0.466 | -0.348 | | | | -0.033 | 0.763 | 0.593 | -0.255 | | | | 0.840 | 0.026 | 0.481 | -0.250 | | |
| la | -0.379 | 0.839 | -0.055 | -0.387 | | | | 0.832 | -0.312 | -0.038 | -0.457 | | | | -0.123 | 0.988 | -0.040 | -0.080 | | |
| ln | -0.505 | -0.750 | -0.186 | -0.384 | | | | -0.823 | -0.331 | -0.077 | -0.455 | | | | -0.879 | -0.164 | -0.006 | -0.448 | | |
| ldmc | 0.656 | 0.052 | -0.747 | -0.093 | | | | 0.017 | 0.699 | -0.701 | -0.141 | | | | 0.836 | -0.053 | -0.495 | -0.231 | | |
| *Fine-root traits* | | | | | | | | | | | | | | | | | | | | |
| **Figure 4d** | **PC1** | **PC2** | **PC3** | **PC4** | | | **Figure 4e** | **PC1** | **PC2** | **PC3** | **PC4** | | | **Figure 4f** | **PC1** | **PC2** | **PC3** | **PC4** | | |
| Variance (%) | 0.521 | 0.320 | 0.140 | 0.019 | | | Variance (%) | 0.494 | 0.360 | 0.134 | 0.012 | | | Variance (%) | 0.506 | 0.315 | 0.152 | 0.027 | | |
| eigenvalue | 1.444 | 1.132 | 0.749 | 0.270 | | | eigenvalue | 1.406 | 1.200 | 0.734 | 0.216 | | | eigenvalue | 1.539 | 0.914 | 0.774 | 0.444 | | |
| SRL | 0.942 | 0.248 | -0.125 | 0.191 | | | | 0.967 | 0.170 | -0.120 | 0.150 | | | | 0.925 | 0.000 | 0.163 | 0.342 | | |
| RDMC | -0.460 | 0.745 | 0.478 | 0.065 | | | | -0.340 | 0.809 | 0.477 | 0.055 | | | | -0.571 | 0.785 | 0.219 | 0.094 | | |
| RD | -0.812 | -0.555 | -0.019 | 0.180 | | | | -0.847 | -0.511 | -0.035 | 0.145 | | | | -0.841 | -0.176 | -0.441 | 0.259 | | |
| RN | 0.572 | -0.596 | 0.563 | -0.007 | | | | 0.457 | -0.704 | 0.544 | -0.006 | | | | 0.690 | 0.435 | -0.574 | -0.066 | | |
| *Leaf and fine-root traits* | | | | | | | | | | | | | | | | | | | | |
| **Figure 4g** | **PC1** | **PC2** | **PC3** | **PC4** | **PC5** | **PC6** | **Figure 4h** | **PC1** | **PC2** | **PC3** | **PC4** | **PC5** | **PC6** | **Figure 4i** | **PC1** | **PC2** | **PC3** | **PC4** | **PC5** | **PC6** |
| Variance (%) | 0.284 | 0.217 | 0.158 | 0.128 | 0.089 | 0.063 | Variance (%) | 0.264 | 0.232 | 0.162 | 0.121 | 0.091 | 0.064 | Variance (%) | 0.395 | 0.218 | 0.138 | 0.106 | 0.059 | 0.038 |
| eigenvalue | 1.508 | 1.317 | 1.126 | 1.013 | 0.842 | 0.712 | eigenvalue | 1.454 | 1.361 | 1.140 | 0.983 | 0.852 | 0.718 | | 1.778 | 1.321 | 1.052 | 0.920 | 0.690 | 0.549 |
| lma | -0.394 | -0.492 | -0.141 | 0.627 | 0.255 | 0.029 | | -0.016 | -0.462 | 0.080 | 0.829 | 0.183 | -0.101 | | -0.632 | -0.518 | -0.118 | 0.377 | 0.298 | 0.226 |
| ldmc | -0.287 | -0.624 | 0.143 | -0.045 | -0.706 | 0.018 | | -0.186 | -0.653 | -0.077 | -0.042 | -0.712 | -0.147 | | -0.769 | -0.355 | -0.045 | -0.281 | 0.261 | -0.343 |
| la | 0.026 | 0.428 | 0.793 | -0.196 | 0.018 | 0.076 | | 0.357 | 0.186 | -0.774 | -0.170 | 0.062 | -0.189 | | -0.033 | 0.191 | 0.893 | 0.376 | 0.118 | -0.047 |
| ln | 0.451 | 0.166 | -0.683 | -0.355 | -0.112 | -0.164 | | -0.365 | 0.318 | 0.700 | -0.222 | 0.028 | -0.350 | | 0.739 | 0.522 | -0.077 | -0.052 | 0.187 | -0.108 |
| SRL | 0.802 | -0.503 | 0.225 | 0.004 | 0.079 | -0.096 | | -0.926 | -0.068 | -0.301 | -0.019 | 0.127 | -0.049 | | 0.685 | -0.613 | 0.143 | 0.124 | 0.015 | -0.253 |
| RDMC | -0.551 | -0.362 | -0.166 | -0.564 | 0.219 | 0.409 | | 0.241 | -0.723 | 0.254 | -0.340 | 0.165 | 0.365 | | -0.329 | 0.508 | -0.444 | 0.618 | -0.023 | -0.215 |
| RD | -0.633 | 0.669 | -0.172 | 0.215 | -0.211 | -0.018 | | 0.848 | 0.363 | 0.208 | 0.174 | -0.224 | -0.043 | | -0.601 | 0.639 | 0.162 | -0.292 | 0.239 | 0.024 |
| RN | 0.696 | 0.242 | -0.155 | 0.319 | -0.185 | 0.545 | | -0.429 | 0.646 | -0.006 | 0.228 | -0.295 | 0.434 | | 0.826 | -0.011 | -0.206 | 0.019 | 0.461 | 0.090 |

**Figure S1.** Changes in mean leaf and fine-root traits between angiosperms and gymnosperms. Significant differences (t-test) within angiosperms and gymnosperms are denoted by * ($P < 0.05$), ** ($P < 0.01$) and *** ($P < 0.001$). Boxes indicate interquartile ranges (the bottom and top parts of the box), and median lines (horizontal lines within boxes) and whiskers indicate the minimum and maximum of the measurement. Abbreviations: ln (measured in % of dry weight), leaf nitrogen concentrations; ldmc (measured in g $g^{-1}$), leaf dry matter content; la (measured in $cm^{-2}$), leaf area; lma (measured in g $cm^{-2}$), leaf mass per area; RN (measured in % of dry weight), root nitrogen concentrations; RD (measured in mm), root diameter; RDMC (measured in g $g^{-1}$), root dry matter content; and SRL (measured in mm $mg^{-1}$), specific root length.

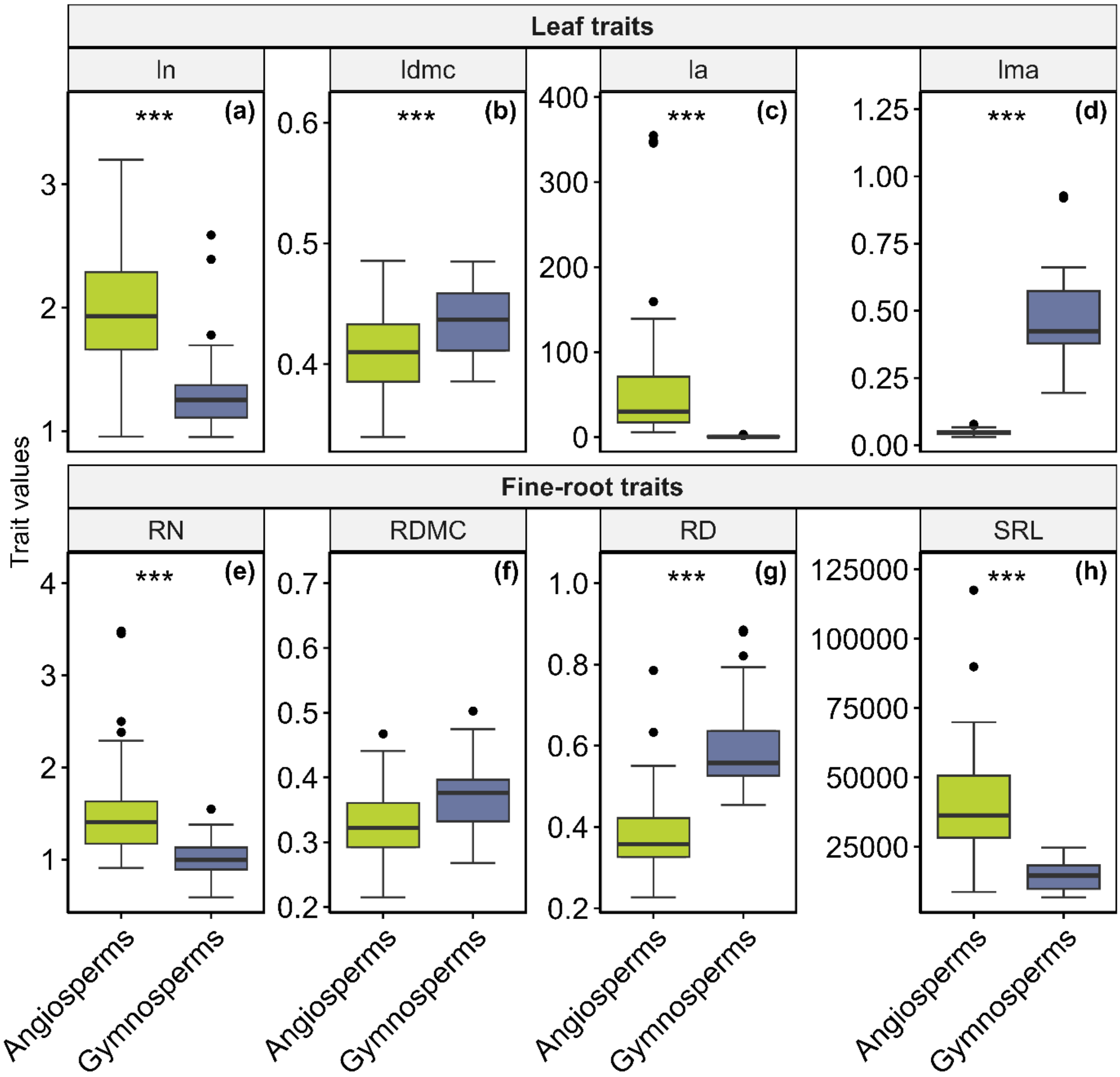

**Figure S2.** Pairwise correlation among leaf and fine-root traits for angiosperm species. The bivariate relationships among leaf and fine-root traits for angiosperm tree species (*n* = 65) are assessed using ordinary least squares (OLS) models and phylogenetic generalized least squares (PGLS) models. Significant correlations are represented by regression lines colored in black for OLS models and in pink for PGLS models. Trait abbreviations follow the caption in Tables S2 and S3.

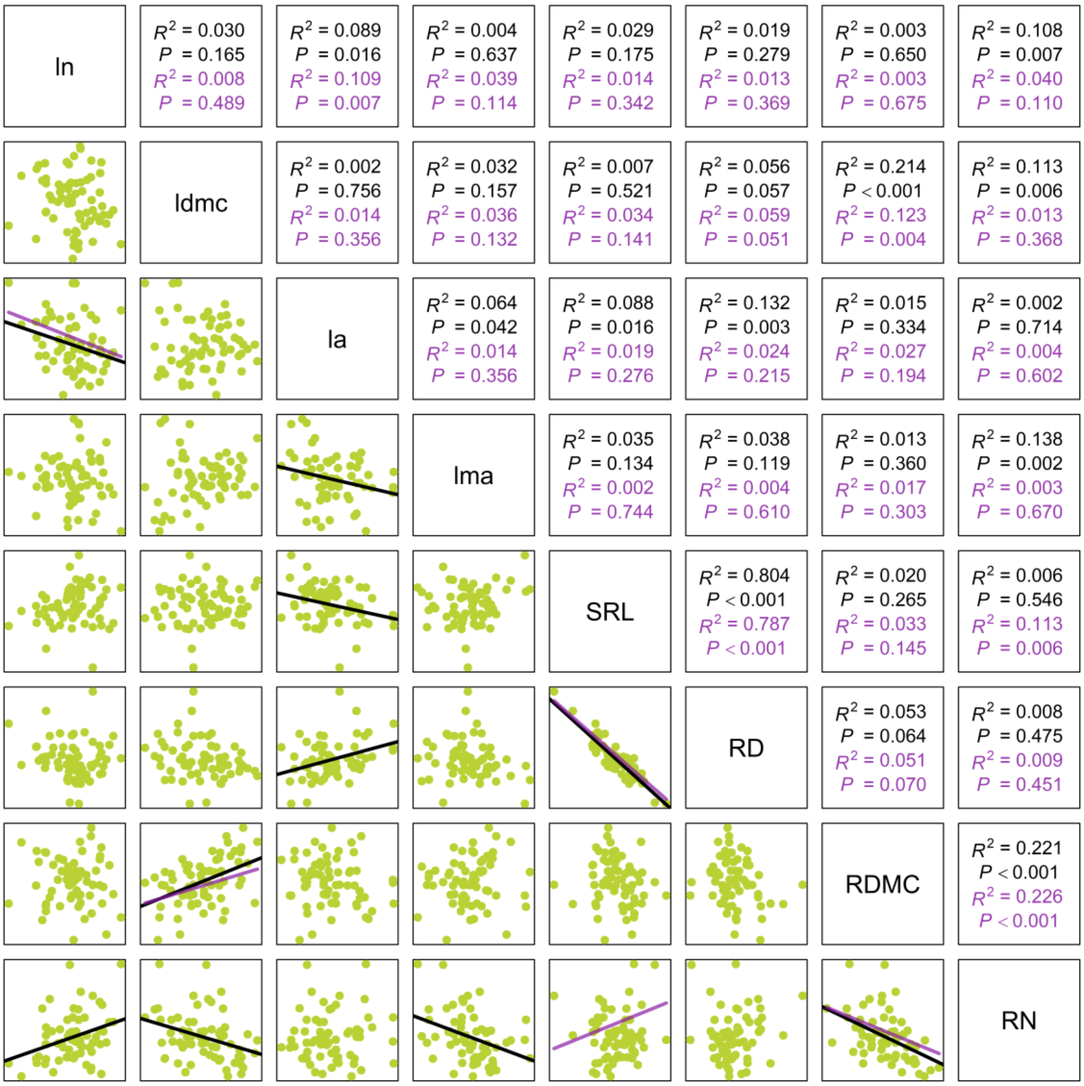

**Figure S3.** Pairwise correlation among leaf and fine-root traits for gymnosperm species. The bivariate relationships among leaf and fine-root traits for gymnosperm tree species (*n* = 25) are assessed using ordinary least squares (OLS) models and phylogenetic generalized least squares (PGLS) models. Significant correlations are represented by regression lines colored in black for OLS models and in pink for PGLS models. Trait abbreviations follow the caption in Tables S2 and S3.

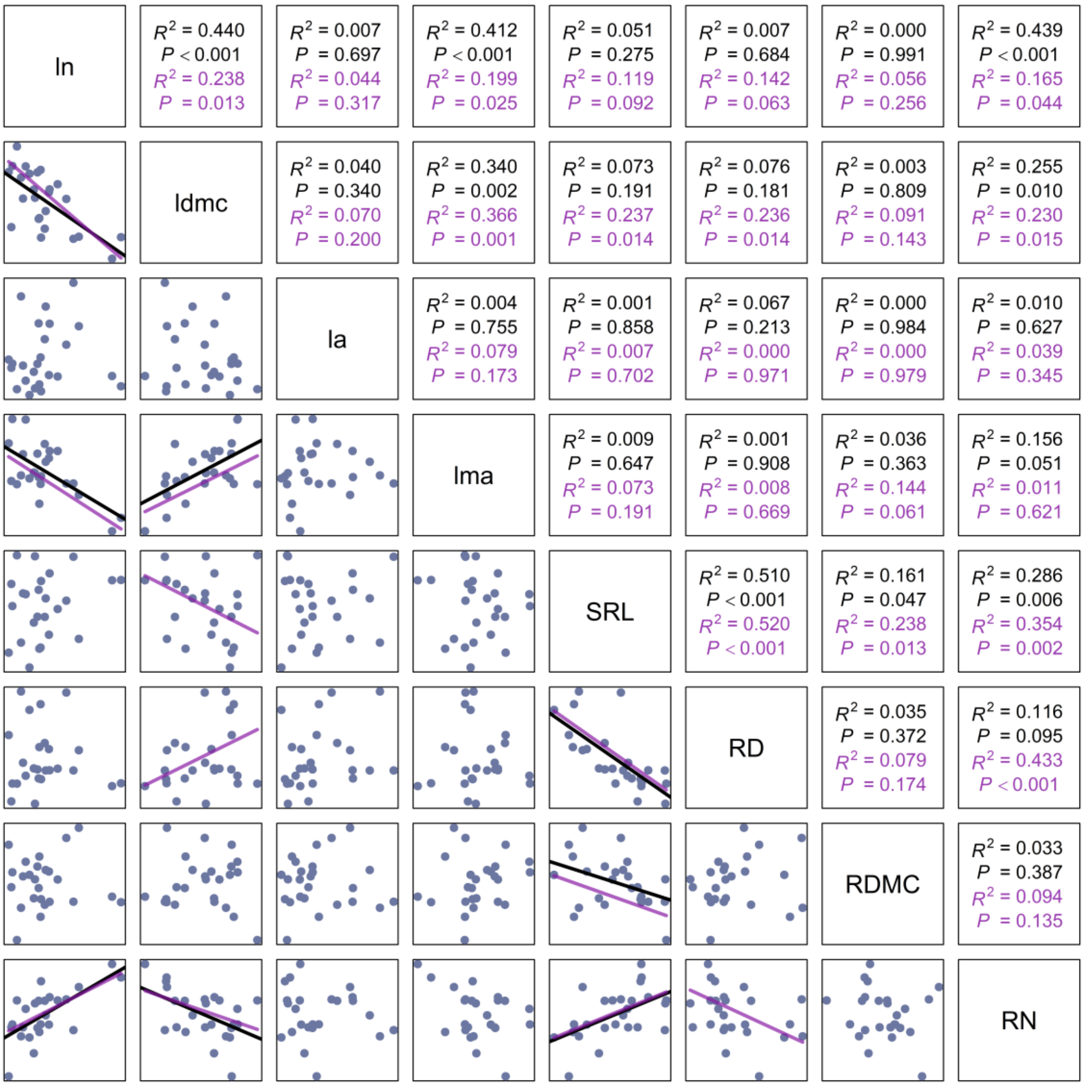

**Figure S4.** Absolute loading scores of phylogenetically informed principal component analyses of the leaf and fine-root traits on the first four principal component (PC1:PC4) axes for all species, angiosperms, and gymnosperms, as shown in Figure 4 and Table S2. lma is leaf mass per area; la is leaf area; ln is leaf nitrogen concentration; ldmc is leaf dry matter content; SRL is specific root length; RDMC is root dry matter content; RD is average root diameter; RN is root nitrogen concentration.

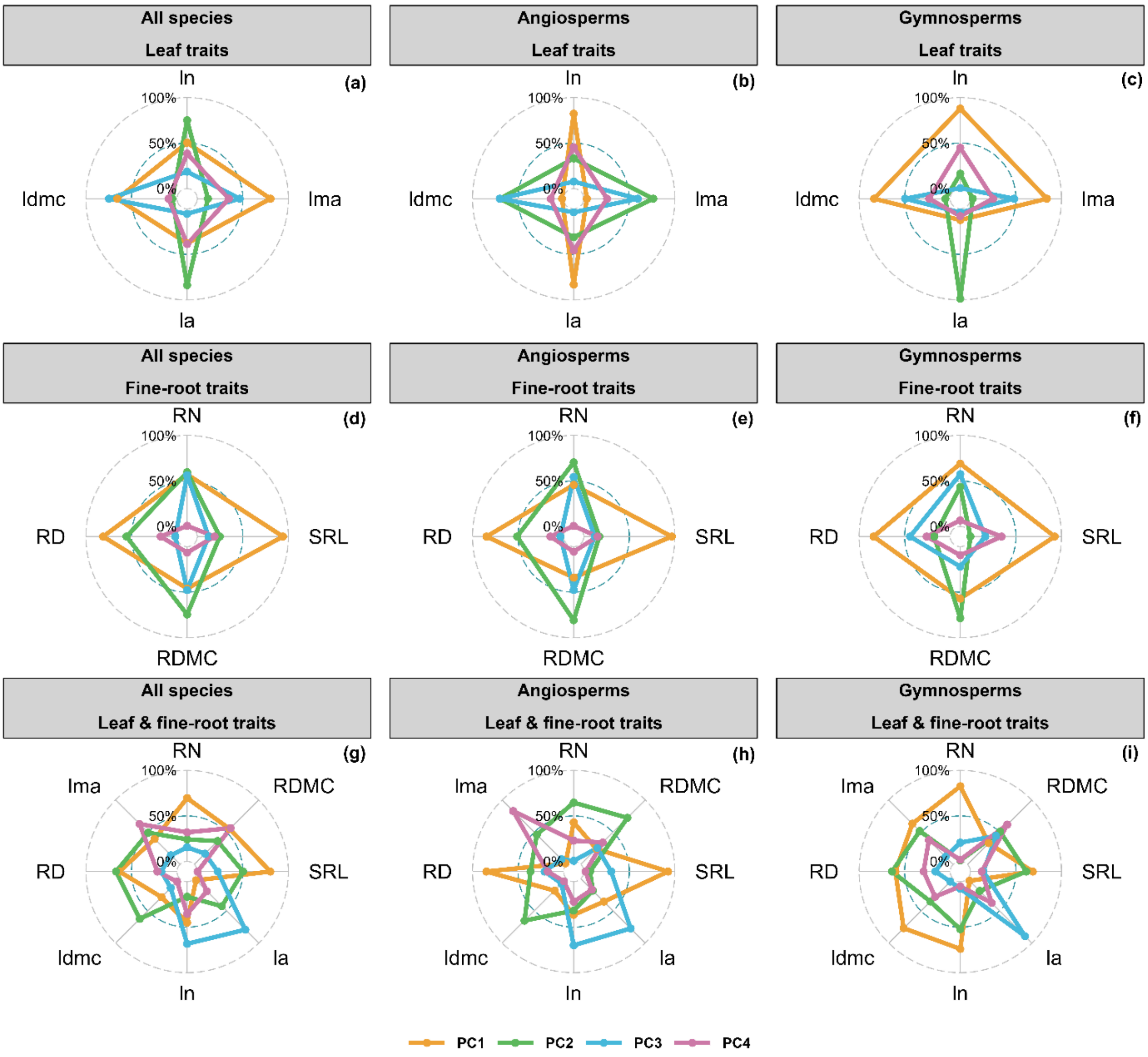
